# Predicting Environmental Chemical Carcinogenicity using a Hybrid Machine-Learning Approach

**DOI:** 10.1101/2021.05.03.442477

**Authors:** Sarita Limbu, Sivanesan Dakshanamurthy

## Abstract

Determining environmental chemical carcinogenicity is an urgent need as humans are increasingly exposed to these chemicals. In this study, we determined the carcinogenicity of wide variety real-life exposure chemicals in large scale. To determine chemical carcinogenicity, we have developed carcinogenicity prediction models based on the hybrid neural network (HNN) architecture. In the HNN model, we included new SMILES feature representation method, by modifying our previous 3D array representation of 1D SMILES simulated by the convolutional neural network (CNN). We used 653 molecular descriptors modeled by feed forward neural network (FFNN), and SMILES as chemical features to train the models. We have developed three types of machine learning models: binary classification models to predict chemical is a carcinogenic or non-carcinogenic, multiclass classification models to predict severity of the chemical carcinogenicity, and regression models to predict median toxic dose of the chemicals. Along with the hybrid neural network (HNN) model that we developed, Random Forest (RF), Bootstrap Aggregating (Bagging) and Adaptive Boosting (AdaBoost) methods were also used for binary and multiclass classification. Regression models were developed using HNN, RF, Support Vector Regressor (SVR), Gradient Boosting (GB), Kernel Ridge (KR), Decision Tree with AdaBoost (DT), KNeighbors (KN), and a consensus method. For binary classification, our HNN model predicted with an average accuracy of 74.33% and an average AUC of 0.806, for multiclass classification, the HNN model predicted with an average accuracy of 50.58% and an average micro-AUC of 0.68, and for regression model, the consensus method achieved R^2^ of 0.40. The predictive performance of our models based on a highly diverse chemicals is comparable to the literature reported models that included the similar and less diverse molecules. Our models can be used in identifying the potentially carcinogenic chemicals for a wide variety of chemical classes.

## INTRODUCTION

Substances capable of causing cancer are known as carcinogens. Carcinogenicity is of primary concern among all the toxicological endpoints due to the severity of its outcome. Carcinogens may be genotoxic that induce DNA damage and induce cancer or non-genotoxic that uses other mode of action such as tumor promotion to exhibit their carcinogenic potential in humans^1^. Some of the genotoxic carcinogens are mutagens too. Many environmental chemicals have been identified as carcinogenic to the humans^2,3^. The onset of cancer in humans depend on various factors including dose and duration of exposure to carcinogens. Identifying carcinogenic compounds is also an integral step during the drug development process. The 2-year rodent carcinogenicity assay has been established since more than 50 years as the standard for the prediction of chemical carcinogenicity^4^. However, such animal testing is time-consuming, costly, and unethical too. The experimentalists need to replace, reduce, and refine (3Rs) the use of animals as this 3Rs policy encourages for alternative methods to minimize the unprincipled use of animals^5^. Computational methods for various toxicological endpoints prediction have now become a popular alternative to the traditional animal testing.

Numerous computational models using machine learning (ML) methods are developed to predict the carcinogenicity based on the properties of chemicals. Computational models can be classification models (qualitative) that predict a chemical is carcinogenic or not (binary classification models) or that predict the degree of carcinogenicity (multiclass classification), and regression models (quantitative) that predict the dose of chemical required for carcinogenesis. Computational models based on structurally related congeneric chemicals are reported to give high predictive performance. Luan et al. reported accuracy of 95.2% while predicting carcinogenicity of N-Nitroso compounds based on support vector machine(SVM) method^6^. Ovidiu Ivanciuc presented SVM based model to predict the carcinogenicity of polycyclic aromatic hydrocarbons (PAH) with 87% accuracy^7^. Computational models based on non-congeneric chemicals are of interest due to their predictive ability for diverse set of chemicals. Fjodorova et al. predicted the carcinogenicity of non-congeneric chemicals with 68% accuracy using counter propagation artificial neural network (CP ANN)^8^. Tanabe et al. reported accuracy of 70% for non-congeneric chemicals based on SVM and improved the accuracy to 80% by developing models on the chemical subgroups based on their structure^9^. Zhang et al. presented binary classification models based on ensemble of eXtreme Gradient Boosting (XGBoost) method that predicted the carcinogenicity of chemicals with 70% accuracy^10^. Li et al. used six different ML methods to generate the binary classification model with 83.91% accuracy and ternary (multiclass) classification models with 80.46% accuracy for the external validation set for the best model^11^. Toma et al. developed binary classification models with accuracy of 76% and 74% and regression models with r^2^ of 0.57 and 0.65 on oral and inhalation slope factors to predict the carcinogenicity for external validation set^12^. Fjodorova et al. reported the correlation coefficient of 0.46 for the test set for their regression models using counter propagation artificial neural network (CP ANN)^8^. Wang et al. constructed a deep learning model that requires less data and achieved 85% accuracy on external validation set for carcinogenicity prediction^16^.

Taken together, numerous carcinogenicity predictive models on congeneric and non-congeneric chemicals for binary classification and a few multiclass and regression models were reported. There is a need for more non-congeneric computational models with a very wide applicability domain for carcinogenicity prediction. In this study, using our hybrid deep learning method and other machine learning methods, we have developed binary classification, multiclass classification and regression models based on very diverse non-congeneric chemicals.

To determine the chemical carcinogenicity, we have developed carcinogenicity prediction models based on the hybrid neural network (HNN) architecture that we reported previously for toxicity prediction^17^. In this study, we used the modified version of the 3D array representation of 1D SMILES to use it in the convolutional neural network (CNN) model^17^. Other machine learning methods used were Random Forest (RF), Bootstrap Aggregating (Bagging) and Adaptive Boosting (AdaBoost) for binary classification and multiclass classification. Random Forest (RF), Support Vector Regressor (SVR), Gradient Boosting (GB), Kernel Ridge (KR), Decision Tree with AdaBoost (DT), and KNeighbors (KN) methods were used for regression.

## MATERIALS & METHODS

### DATA

We have collected carcinogens from several different data sources detailed below.

1. Chemical Exposure Guidelines for Deployed Military Personnel Version 1.3 (MEG). We curated the carcinogenic chemicals from the Technical Guide 230 (TG230) in the “Chemical Exposure Guidelines for Deployed Military Personnel”^18^. TG 230 provides military exposure guidelines (MEGs) for chemicals in air, water, and soil along with an assigned carcinogenicity group for each chemical. Chemicals are categorized into one of the 5 groups: Group A (Human Carcinogen), Group B (Probable human carcinogen), Group C (Possible human carcinogen), Group D (Not classifiable) and Group E (No evidence of carcinogenicity).
2. Environmental Health Risk Assessment and Chemical Exposure Guidelines for Deployed Military Personnel 2013 Revision (TG230). We curated the carcinogenic chemicals listed in the Technical Guide 230 (TG230) in the “Environmental Health Risk Assessment and Chemical Exposure Guidelines for Deployed Military Personnel”^19^ provides with military exposure guidelines (MEGs).
3. National Toxicology Program (NTP). Carcinogenic chemicals were curated from the NTP^20^. NTP lists two groups of carcinogenic chemicals: 1) reasonably anticipated to be a human carcinogen and 2) known to be human carcinogens.
4. International Agency for Research on Cancer (IARC) Carcinogenic chemicals were curated from IARC^21^. IARC categorizes chemicals into one of the 5 groups: Group 1 (Carcinogenic to humans), Group 2A (Probably carcinogenic to humans), Group 2B (Possibly carcinogenic to humans), Group 3 (Not classifiable as to its carcinogenicity to humans), and Group 4 (Probably not carcinogenic to humans).
5. The Japan Society for Occupational Health (JSOH) Carcinogenic chemicals were curated with recommendation of Occupational Exposure Limits published by the JSOH^22^ which are classified into one of the 3 groups: Group 1 (carcinogenic to humans), Group 2A (probably carcinogenic to humans), and Group 2B (possibly carcinogenic to humans).
6. The National Institute for Occupational Safety and Health (NIOSH) Carcinogenic chemicals curated from the NIOSH^23^.
7. Carcinogenic Potency Database (CPDB)
  a. CPDB_CPE (CPDB CarcinoPred-EL) data: CPDB data for rat carcinogenicity collected from the CarcinoPred-EL developed by Zhang et al^10^. The list contains 494 carcinogenic and 509 non-carcinogenic chemicals.
  b. CPDB data: CPDB^24^ data were collected and processed to obtain the median toxicity dose (TD50) for rat carcinogenicity. TD50 is the dose-rate in mg/kg body wt/day administered throughout life that induces cancer in half of the test animals. 561 carcinogenic chemicals were obtained with TD50 values for rat carcinogenicity. 605 noncarcinogenic chemicals were obtained for rat carcinogenicity. For 543 carcinogenic chemicals out of 561, the TD50 values in mmol/kg body wt/day were also obtained from DSSTox database (https://www.epa.gov/chemical-research/distributed-structure-searchable-toxicity-dsstox-database).
8. Chemical Carcinogenesis Research Information System (CCRIS). Carcinogenesis data were collected from the CCRIS at ftp://ftp.nlm.nih.gov/nlmdata/.ccrislease/. The carcinogenicity and mutagenicity data were extracted. 6,833 chemicals were obtained after eliminating duplicates/conflicting data when compared to data sources 1 to 6, out of which 4,054 were carcinogenic/mutagenic and 2,779 were non-carcinogenic/mutagenic.
9. Drugbank 2018 The drugs data were collected from the drugbank (www.drugbank.ca). The approved drugs predicted as carcinogenic by Zhang et al^10^ were removed, remining 1,756 approved drugs were considered as non-carcinogenic.

#### Dataset I: Binary Classification Data

The two classes considered in the binary classification models were class 0 (non-carcinogen) and class 1 (carcinogen). Datasets used to train the models are listed below:

i. For binary classification of chemicals to predict carcinogenic or non-carcinogenic category, 448 carcinogenic chemicals were obtained from data sources 1 to 6.
  Data 1 (MEG): The chemicals classified into Groups A, B and C were considered as carcinogens.
  Data 2 (TG30): The chemicals listed as carcinogens were considered as carcinogens.
  Data 3 (NTP): The chemicals classified as either “reasonably anticipated to be a human carcinogen” or “known to be human carcinogens” were considered as carcinogens.
  Data 4 (IARC): The chemicals classified into Groups 1, 2A, and 2B were considered as carcinogens.
  Data 5 (JSOH): The chemicals classified into Groups 1, 2A, and 2B were considered as carcinogens.
  Data 6 (NIOSH): The carcinogenic chemicals listed were considered as carcinogens.
ii. CPDB_CPE chemicals from data source 7a contributed 320 carcinogenic and 458 non-carcinogenic additional data after comparing to the data from data sources 1 to 6 and removing duplicates and conflicting chemicals.
iii. The CCRIS mutagenicity/carcinogenicity data from data source 8 contributed 3,868 mutagenic/carcinogenic data and 2,500 non-mutagenic/carcinogenic data.
iv. 400 non-carcinogenic approved drugs from data source 9 were also used in this classification model.

The dataset I for the binary classification model, we used a total of 7,994 chemicals with 4,636 carcinogenic and 3,358 noncarcinogenic chemicals.

#### Dataset II: Multiclass Classification Data

The classes considered in the multiclass classification models were class 0 (non-carcinogen), 1 (possibly carcinogen and not classifiable chemicals) and 2 (carcinogen and probably carcinogen). Datasets used to train the models listed below:

i. For multiclass classification, 882 carcinogenic chemicals and 2 non-carcinogenic chemicals were collected from data sources 1,3,4, and 5. There were total of 2 in class 0, 604 in class 1, and 278 in class 2 in this dataset.
  Data 1 (MEG): The chemicals classified into Groups A, B were considered as class 2. The chemicals classified into Groups C and D were considered as class 1 carcinogens. Chemicals classified into group E are considered as class 0 compounds.
  Data 3 (NTP): The chemicals classified as either “reasonably anticipated to be a human carcinogen” or “known to be human carcinogens” were considered as class 2.
  Data 4 (IARC): The chemicals classified into Groups 1 and 2A were considered as class 2 carcinogens and those classified into Groups 2B and 3 were considered as class 1 carcinogens.
  Data 5 (JSOH): The chemicals classified into Groups 1 and 2A were considered as class 2 carcinogens and those classified into Groups 2B were considered as class 1 carcinogens.
  Considering Group D of MEG data as class1 carcinogen along with Group C and considering Group 3 of IARC data as class 1 carcinogen along with Group 2B increased the multiclass data significantly in this dataset. In the case of binary classification, we discarded these groups.
ii. CPDB chemicals from data source 7b contributed 277 carcinogenic and 457 non-carcinogenic additional data after removing duplicates and conflicting chemicals compared to the data from data sources 1, 3, 4 and 5. The 277 carcinogenic chemicals were categorized into class 2 and 457 noncarcinogenic chemicals were categorized into class 0.

The dataset II for the multiclass classification models we used a total of 459 chemicals data in class 0, 604 chemicals data in class 1 and 555 chemicals data in class 2.

#### Dataset III: Regression Data

Regression models were developed to predict the quantitative carcinogenicity or the Median Toxic Dose (TD50) of the chemicals in the form of pTD50 (logarithm of inverse of TD50). Dataset III for the regression models consisted of 561 TD50 data in mg/kg body wt/day converted to pTD50 from data source 7b. Independently, the regression models were also developed on 543 TD50 data in mmol/kg body wt/day converted to pTD50.

### Descriptors

Mordred descriptor calculator^25^ that calculates 1,613 2D molecular descriptors from SMILES is used for descriptor calculation. This descriptor calculator supports Python 3. The final set of 653 descriptors were obtained with no missing calculated values for the entire datasets for which descriptors were calculated. The 653 descriptors were used as a final set of features for the training and test data in the machine learning models.

### SMILES Preprocessing

The simplified molecular-input line-entry system (SMILES) uses ASCII strings for the 1D chemical structure representation of compound and can be used to convert to its 2-D or 3-D representation. It is one of the key chemical attributes and is used in our deep learning model. Raw texts cannot be directly used as input for the deep learning models but should be encoded as numbers. Tokenizer class in python is used to encode the SMILES string. The SMILES preprocessing method that we used while predicting toxicity^17^ created the index for the set of unique characters of SMILES from the training set only. Here, we have created unique index for 94 characters in the ASCII table so that there is no possibility of missing out creating index of any character in the SMILES string represented in any format. 94 characters from ‘!’ to ‘~’ in the ASCII table represented by decimal numbers 33 to 126 made the vocabulary of the possible characters in the SMILES. A dictionary D ={‘!’:1, ‘”’:2, ‘#’:3, ‘$’:4, …, ‘C’:35, …, ‘~’:94} is created that maps every character in the list of 94 ASCII characters to a unique index.

Each character in the SMILES is converted to its corresponding index in the dictionary D and a vector is created for the SMILES of each compound. As an example, acrylonitrile-d3 with SMILES string C=CC#N is encoded as [35, 29, 35, 35, 3, 46]. As the length of the SMILES vary depending on the length and properties of the compound, the length of the encoding results also varies. The resulting vector for the SMILES of every input compound is thus padded with 0s or truncated so that they are of uniform length, L. The SMILES for the input compounds are converted to a 2-D matrix of size K x L where K is the number of input SMILES and L = 325 is the allowed maximum length of the SMILES string used in the model. Our previous method^17^ mapped the SMILES for the K number of chemicals to a one-hot encoded matrix of size KxLxM where M is the number of the possible characters in the SMILES.

### Machine Learning Models

#### Hybrid Neural Network Model

Hybrid Neural Network (HNN) model^17^ that we developed for chemical toxicity prediction was used here by modifying the SMILES vectorization method and then the method by which the vectorized SMILES input is processed by Convolutional Neural Network (CNN) of the model. The model is developed in python using the Keras API with Tensorflow in the backend. The model consists of a CNN for deep learning based on structure attribute (SMILES) and multilayer perceptron (MLP) type feed forward neural network (FFNN) for learning based on descriptors of the chemicals. To vectorize SMILES, each character in the SMILES string is converted to its positional index in the dictionary as explained in the SMILES preprocessing section. The 2D array of vectorized SMILES strings were the input for the CNN. Embedding layer of Keras is used to convert index of each character in the SMILES string into a dense vector. The embedding layer takes 3 arguments as input: input_dim is the vocabulary size of the characters in the SMILES string, output_dim is the size of the embedded output for each character, and input_length is the length of the SMILES string. In the model, we have embedded index of each character in the SMILES to a vector of size 100 by setting the output_dim to 100. The embedding layer converts the input 2D array of size KxL, where K is the number of SMILES and L is the maximum length of SMILES, to 3D array of size KxLx100.

1D convolution layer Activation function ReLU represented mathematically as max(0, x) is used in the model that replaces all the negative values with zeros. The derivative of ReLU is always 1 for positive input that counteracts the vanishing gradient problem during the backpropagation. The output of the pooling layer of the CNN, together with the FFNN are connected to the final fully connected layer to perform the classification task.

#### Other Machine learning Algorithms

In the case of binary classification and multiclass classification, to test the performance of HNN, the other machine learning algorithms were used were Random Forest (RF), Bootstrap Aggregating (Bagging) using Bagged Decision Tree, and Adaptive Boosting (AdaBoost).

Random Forest (RF): is a bootstrap aggregating (bagging) model and uses ensemble of decision trees for making final decision. This algorithm uses only a subset of features to find the best feature in order to separate classes at each node of the tree.

Bagged Decision Tree (Bagging): uses bootstrap method to reduce variance and overfitting. It uses ensemble method for the final decision. Bagging method uses all features to find the best feature for splitting node of the tree.

Adaptive Boosting (AdaBoost): is also an ensemble method of machine learning that uses weak classifiers to make stronger classifiers.

Several regression models were developed based on Random Forest (RF), Support Vector Regressor (SVR), Gradient Boosting (GB), Kernel Ridge (KR), Decision Tree with AdaBoost (DT), and KNeighbors (KN) using the sklearn package in python to make the final consensus prediction of the median toxic dose (TD50). A consensus prediction based on average of all seven predicted values was calculated.

Random Forest (RF): is a bootstrap aggregating (bagging) model and uses ensemble of decision trees for making final prediction. The regression model is fit for every feature and the data is split at several points. The feature with least error is selected as the node.

Support Vector Regressor (SVR): SVR depends on the subset of training data. SVR performs non-linear regression using kernel trick and transforms inputs into m-dimensional feature space.

Gradient Boosting (GB): it produces ensemble of weak prediction model or regression trees in a stage-wise fashion. In each stage, it optimizes a loss function by choosing the function that point in the negative gradient direction.

Kernel Ridge (KR): Ridge regression uses L2 regularization to limit the size of the coefficients of the model and eliminates the problem in least square regression. Ridge method adds penalty to the coefficients equal to the square of the magnitude of coefficients. Regularization parameter λ controls the penalty term. Kernel ridge uses kernel trick to make the model non-linear.

Decision Tree with AdaBoost (DT): The prediction of decision tree was boosted with AdaBoost. Decision Tree method makes the prediction by learning decision rules from the training data. AdaBoost is a boosting algorithm introduced by Freund and Schapire^26^. AdaBoost makes final predictions from weighted voting of the individual predictions from weak learners. It implements AdaBoost.R2 algorithm^27^.

Kneighbors (KN): Nearest neighbors finds k number of training data closest to the test data for which prediction is done. Each of the closest neighbors contributes equally while making prediction (default parameter).

#### Model Evaluation

All the results presented for the model evaluation are the average of 10 repeats (in case of binary classification models and regression models) and 30 repeats (in case of multiclass classification models). Approximately 20% of data were separated randomly in each iteration as test set and remaining data as training set like 5-fold cross validation except that the test sets were randomly selected in each iteration.

In case of binary classification and multiclass classification, the performance of each model was evaluated based on accuracy and Area Under the receiver operating characteristic Curve (AUC). The classification models were also evaluated for the sensitivity, and specificity. The evaluation scores are calculated as:

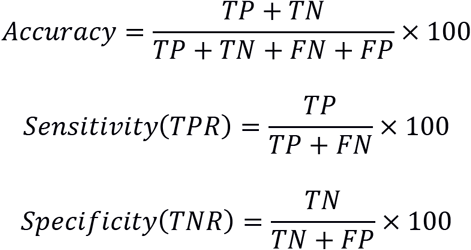

In the multiclass classification, micro averaging is used to obtain the average of the metrics of all the classes. Micro averaging involves calculating the average by converting the data in multiple classes to binary classes and giving equal weight to each observation. In multiclass classification with imbalanced dataset, micro averaging of any metric is preferred when compared to macro averaging which involves calculating the metrics separately for each class and then averaging them by giving equal weight to each class. In case of multiclass classification with *n* number of classes,

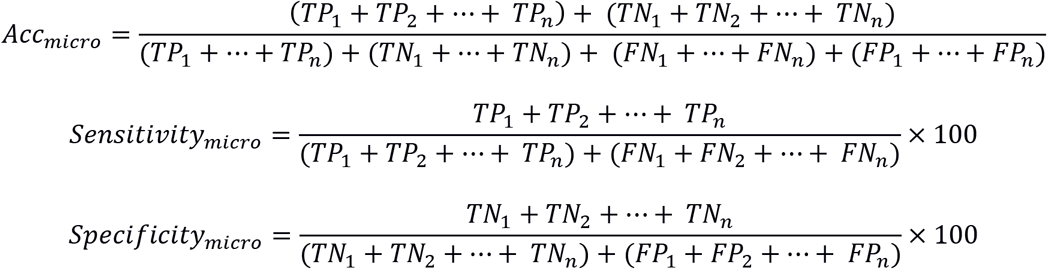

Where, TP = True Positive, TN = True Negative, FP = False Positive, FN = False Negative, TPR = True Positive Rate, TNR = True Negative Rate.

The performance of each regression models was evaluated based on Coefficient of Determination (R^2^). The coefficient of determination gives the percentage of variation in the dependent variable that is predictable from the independent variable or that is explained by the independent variable.

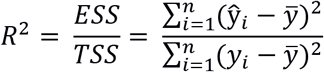

Where, ESS is explained sum of squares and TSS is the total sum of squares, 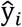 is the predicted value of the *i^th^* dependent variable, *y_i_* is the *i^th^* observed dependent variable, and 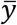 is the mean of the observed data.

## RESULTS AND DISCUSSION

To predict chemical carcinogenicity, binary and multiclass classification models were developed based on the HNN, RF, Bagging, and AdaBoost methods. Regression models were developed based on the HNN, RF, SVR, GB, KR, DT and KN methods to predict the Median Toxic Dose (TD50) of the chemicals. In the HNN models, we used the modified version of the 3D array representation of 1D SMILES to use it in the convolutional neural network (CNN) from our previous model^17^. The SMILES processing method here included a vocabulary of 94 characters in the ASCII table not to miss any possible characters of SMILES in any format. Also, instead of using one-hot encoding to vectorize the characters in the 1-D SMILES, embedding layer of the CNN was used.

### Carcinogenicity Prediction using Binary Classification

The binary classification models were developed for the Dataset I comprising of 7,994 chemicals (4,636 carcinogenic and 3,358 noncarcinogenic) from 9 different sources. Out of 1,613 descriptors calculated by Mordred descriptor calculator, 653 descriptors with no missing values were used to develop the models. In the HNN model we used the SMILES string in addition to the 653 descriptors. The average accuracy, AUC, sensitivity, and specificity of the HNN, RF, and Bagging models were comparable whereas AdaBoost statistical metrics were significantly lower (Figure 1). The average accuracy of the three models was 74% and their average AUC was ~0.81. The average sensitivity and specificity of HNN model were 79.47% and 67.3%.

**Figure 1.**
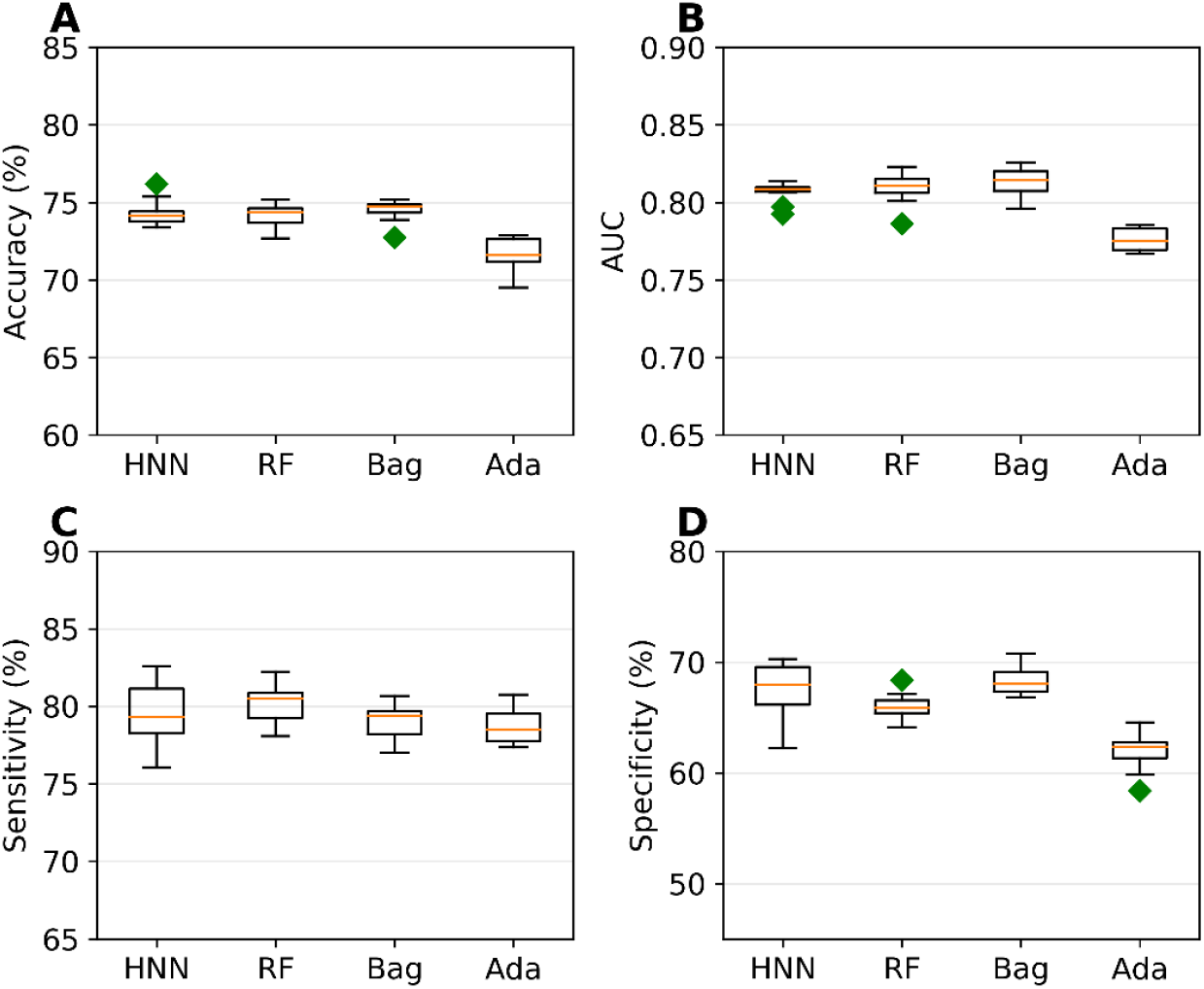
A) Accuracy, B) AUC, C) Sensitivity, and D) Specificity for the dataset I as given by the binary classification models based on the HNN, RF, Bagging, and AdaBoost

Zhang et al.^10^ also built several machine learning models on the CPDB’s 1003 carcinogenic data on rats and the highest average accuracy they reported was 70.1% and AUC of 0.765 for the five-fold cross-validation. Wang et al. developed a novel deep learning tool CapsCarcino on the 1003 rat data from CPDB used by Zhang et al. For five-fold cross-validation they reported average accuracy of 74.5%, sensitivity of 75%, and specificity of 74.2%. Li et al. developed 30 models on only 829 rat data from CPDB with highest average accuracy of 89.29% on their test set. Tanabe et al. developed SVM model with an accuracy of 68.8% and AUC of 0.683 for non-congeneric chemicals from 6 sources using dual cross-validation and they improved the accuracy by developing models on congeneric subgroups. These studies clearly demonstrate that models developed on more diverse chemicals results in reduced accuracy. The predictive performance of our models based on a highly diverse set of chemicals is still good in comparison to the earlier models with a high AUC. Hence, we expect our carcinogenicity predicting models to make carcinogenicity prediction for a wider variety of chemicals.

### Carcinogenicity Prediction using Multiclass Classification

The multiclass classification models were developed for the Dataset II contains 1,618 chemicals with 459 chemicals in class 0, 604 chemicals in class 1 and 555 chemicals in class 2 whereas class 0 comprises of non-carcinogens, class 1 comprises of possible carcinogens and not classifiable chemicals, and class 2 comprises of carcinogens and probable carcinogens. The average overall accuracy is 50.58%, 54.73 %, 55.52%, and 46.50%, the average micro accuracy is 67.05%, 69.82%, 70.34% and 64.33% whereas the average micro AUC is .68, .724, .725, and .653 for HNN, RF, Bagging, and AdaBoost respectively (Figure 2). As observed by Limbu et al^17^, the HNN model is not performing better in comparison to RF and Bagging method. This is possibly because deep learning method performs best with a large dataset and the dataset used in these two studies are not sufficiently large enough.

**Figure 2.**
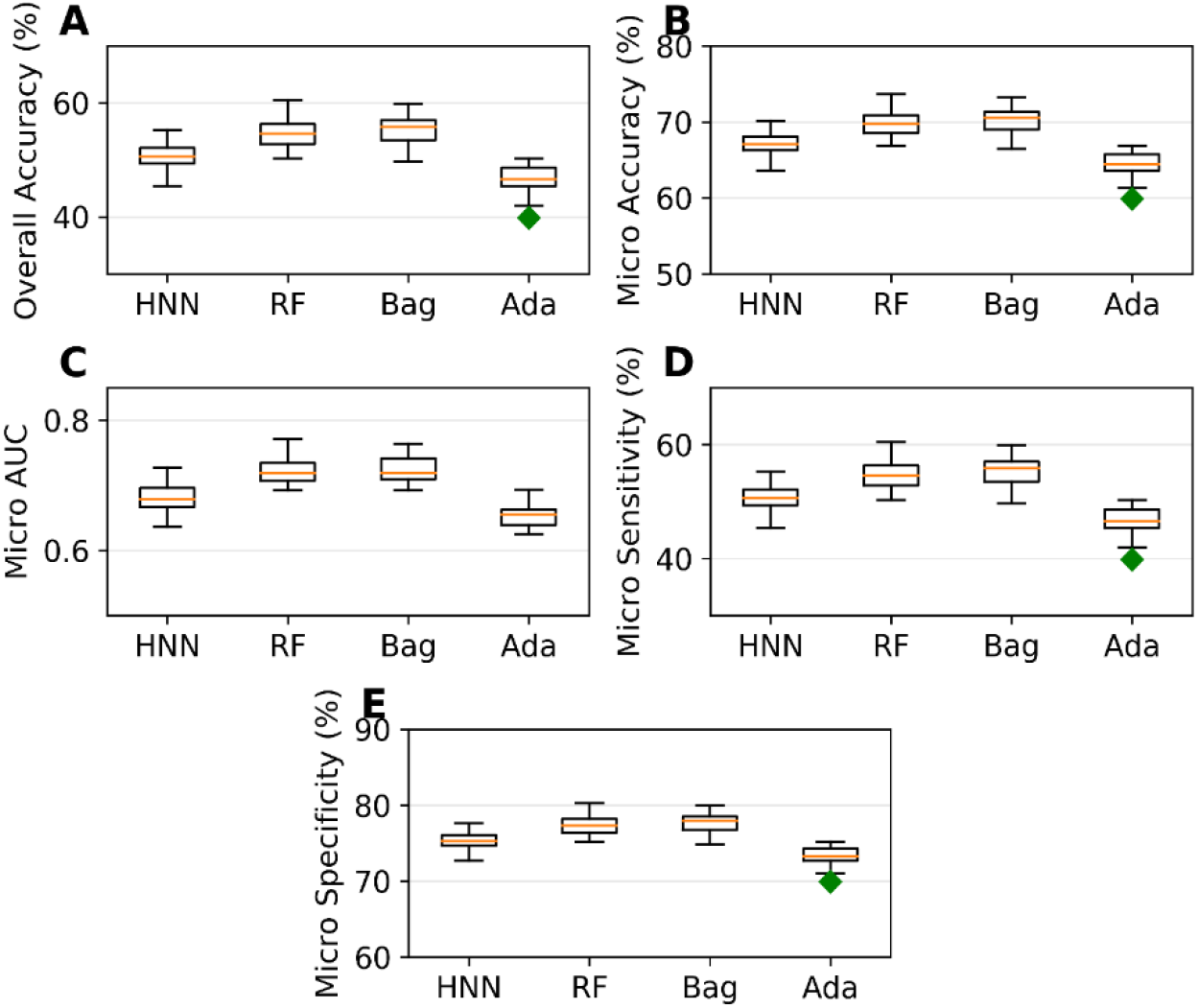
A) Overall accuracy, B) micro accuracy, C) micro AUC, D) micro sensitivity, and E) micro specificity for the dataset II as given by the multiclass classification models based on HNN, RF, Bagging, and AdaBoost methods.

Li et al. developed 30 multiclass (ternary) classification models that categorized compounds into carcinogenic I (strongly carcinogenic), carcinogenic II (weakly carcinogenic) and non-carcinogens^11^. The kNN model based on MACCS fingerprint with best predictive performance achieved average micro accuracy of 81.89%. The ternary classification of their data was based on the TD50 values where TD50 ≤ 10 mg/kg/day were carcinogenic I and TD50 > 10 mg/kg/day were carcinogenic II whereas classification of data in our models are based on their category, they are class 2 if they are carcinogenic or probably carcinogenic, class 1 if they are possibly carcinogenic or not classifiable chemicals, class 0 if they are non-carcinogenic. All the data from CPDB with TD50 were classified as class 2, and non-carcinogens were classified as class 0 but none of them as class 1.

### Carcinogenicity Prediction using Regression

Regression models were developed for the Dataset III comprising of 561 TD50 chemicals. The models predicted the carcinogenicity in the form of pTD50 (logarithm of inverse of TD50) and the average of all the 7 predicted values was calculated as the final consensus prediction of pTD50 value. The average R^2^ is 0.35, 0.36, 0.04, 0.33, 0.36, 0.39, and 0.21 for the HNN, RF, SVM, GB, KR, DTBoost, and NN methods respectively (Figure 3). The overall R^2^ was slightly increased to 0.40 by the consensus prediction. The average correlation coefficient (R) is 0.628, 0.611, 0.322, 0.588, 0.614, 0.636, 0.527, and 0.649 for the HNN, RF, SVM, GB, KR, DTBoost, NN, and consensus methods respectively (Figure 4). The models were also developed for 543 TD50 data in mmol/kg body wt/day. The average correlation coefficient (R) is 0.604, 0.601, 0.287, 0.577, 0.545, 0.617, 0.497, and 0.629 for the HNN, RF, SVM, GB, KR, DTBoost, NN, and consensus methods respectively (Figure 5).

**Figure 3.**
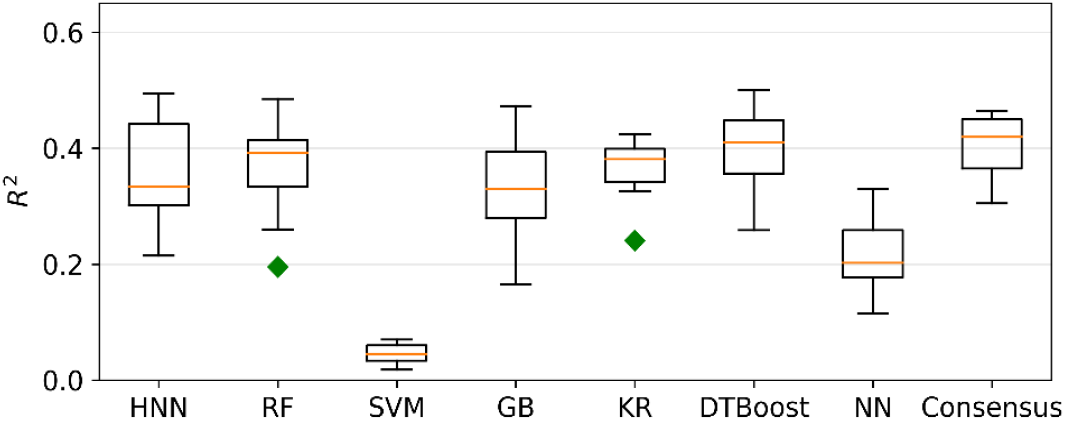
R^2^ of regression models based on HNN, RF, SVM, GB, KR, DTBoost, NN, and Consensus methods.

**Figure 4.**
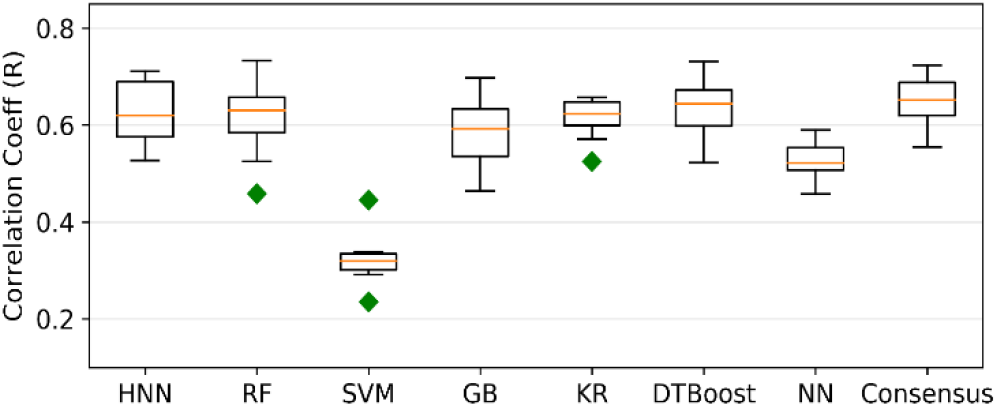
Correlation coefficient (R) of regression models based on HNN, RF, SVM, GB, KR, DTBoost, NN, and Consensus methods.

**Figure 5.**
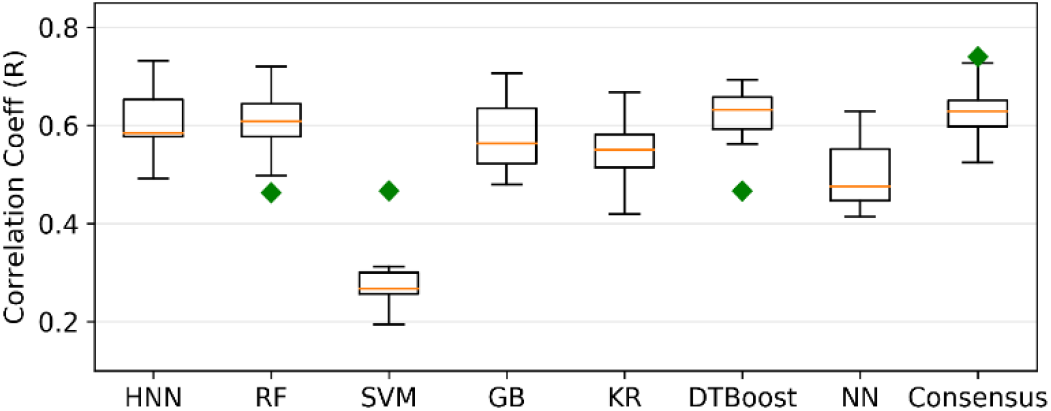
Correlation coefficient (R) of regression models based on HNN, RF, SVM, GB, KR, DTBoost, NN, and Consensus methods that predicts the carcinogenicity in mmol/kg body wt/day.

Fjodorova et al. developed the quantitative models for carcinogenicity prediction on 805 rat data from CPDB using counter propagation artificial neural network (CP ANN)^8^. The correlation coefficient of the models was 0.46 for the test set. Toma et al. developed regression models to predict the carcinogenicity for external validation set with r^2^ of 0.57 and 0.65 for models using oral and inhalation slope factor^12^. In this study, only 315 out of 1110 oral compound and 263 out of 990 inhalation compounds were included in their final dataset after selecting compounds based on various criteria. The external validation set was randomly selected from the finally obtained dataset with highly similar compounds. This may the reason for a slightly better coefficient of determination reported by Toma et al in comparison to our models. Singh et al. developed regression models based on generalized regression neural network (GRNN) to predict the carcinogenicity in mmol/kg body wt/day for 457 CPDB compounds and reported correlation coefficient 0.896 ^28^. The high value of correlation coefficient in comparison to our models could be attributed to the nine molecular descriptors selected for the regression models and the GRNN method was used.

## CONCLUSIONS

Determining environmental chemical carcinogenicity is an urgent need. Though several machine learning models have been reported, there is a need for more non-congeneric computational models with a very wide applicability domain for carcinogenicity prediction. In this study, we determined the carcinogenicity of thousands wide variety class of real-life exposure chemicals. To determine chemical carcinogenicity, we have developed carcinogenicity prediction models based on our hybrid neural network (HNN) architecture. In the HNN model, we included new SMILES feature representation method. Using our hybrid deep learning method and other machine learning methods, we have developed binary classification, multiclass classification and regression models based on very diverse non-congeneric chemicals. The binary and multiclass classification models developed for the larger set of diverse chemicals were from a diverse source, and most of them chemicals were human exposure relevant chemicals.

To compare our model performance, we have developed models based on the other machine learning methods such RF, Bagging, SVM and Ada boost. The models based on the HNN, RF and Bagging methods predicted the carcinogens with an average accuracy of 74% and average AUC of 0.81 which shows that the carcinogen predictions made by these models can be considered highly reliable. Multiclass classification models to categorize the carcinogenicity of chemicals into one of the 3 class: non-carcinogens, possible carcinogens/not classifiable chemicals, or carcinogens/probable carcinogens, were developed. The HNN model exhibited an average accuracy of 50.58%, an average micro accuracy of 67.05%, and average micro AUC of .68. We also developed the regression models to predict the median toxic dose of chemicals in the form of pTD50. The overall R^2^ of 0.40 was achieved by the consensus prediction by calculating the average of all the methods. Though our model included with the very diverse chemical categories and larger number of chemicals from different data sources, still our models could be able to predict the binary, categorical (multiclass), and quantitative (regression) carcinogenicity comparable to the other literature reported models that included with smaller and similar chemicals. Therefore, our models can be used to identify the potential carcinogens for any type of chemicals.

## ACKNOWLEDGMENTS

The author Sivanesan Dakshanamurthy (SD) wishes to acknowledge the support in part by the United States Department of Defense (DOD) grant CA140882. The author Sivanesan Dakshanamurthy wishes to acknowledge the support by the GUMC Lombardi Comprehensive Cancer Center, the CCSG grant P30 CA051008/CA/NCI NIH HHS/United States, and the GUMC Computational Chemistry Shared Resources (CCSR).

